# Multi-Target In Silico Investigation of Withaferin A as a Potential Antiviral Inhibitor Against Key Marburg Virus Proteins

**DOI:** 10.64898/2026.03.06.710011

**Authors:** Farayed Ahamed Nabil, Abu Darda, Ehsanul Islam, Ferdaus Mohd Altaf Hossain, Kazi Md. Ali Zinnah

**Affiliations:** Faculty of Biotechnology and Genetic Engineering, Sylhet Agricultural University, Sylhet-3100, Bangladesh; Department of Animal and Fish Biotechnology, Sylhet Agricultural University, Sylhet-3100, Bangladesh; Department of Pharmacy, Faculty of Biological Science, Jahangirnagar University, Savar, Dhaka-1342, Bangladesh; Department of Dairy science, Sylhet Agricultural University, Sylhet-3100, Bangladesh

**Keywords:** Marburg virus, Withaferin A, VP35, Nucleoprotein, Molecular docking, Structure-based drug discovery

## Abstract

Marburg virus (MARV) is a highly pathogenic filovirus that causes hemorrhagic fever with a high mortality rate, with very limited treatment options. The urgent need for targeted antiviral agents emphasizes the importance of structure-based drug discovery approaches. The present study aimed to evaluate the antiviral potential of Withaferin A (PubChem CID-265237) against three key proteins of MARV: viral protein 35 (VP35), and nucleoproteins (NP). Three-dimensional structures of these proteins were retrieved from RCSB-Protein Data Bank and docked with Withaferin A using AutoDock Vina. The ligand demonstrated favourable binding affinities towards all three viral targets, indicating strong interaction potential at functionally relevant sites. Drug-likeness and pharmacokinetic properties predicted using SwissADME and pkCSM indicated acceptable ADMET profiles that comply with key drug-like criteria. To validate the stability of the docking, molecular dynamics simulations (GROMACS, 100 nanoseconds) were conducted. The protein-ligand complexes exhibited stable root mean square deviation (RMSD), root mean square fluctuation (RMSF), and consistent hydrogen bonding patterns throughout the simulation. The MM-GBSA binding free energy analysis further supported favorable binding energetics, predominantly driven by van der Waals and electrostatic interactions. Altogether, these findings demonstrate that Withaferin A exhibits promising multi-target inhibitory potential against key MARV proteins. This study provides molecular insights into ligand-protein interactions and supports further experimental validation of Withaferin A as a potential therapeutic candidate against Marburg virus.

## 1. Introduction

Marburg virus (MARV) is an extremely pathogenic filamentous, negative (negative sense) single-stranded RNA virus belonging to the Filoviridae family, the same virus family as Ebola. It causes Marburg virus disease (MVD), a severe hemorrhagic fever with high fatality rates, systemic inflammation, multiorgan failure, and immune dysregulation (1). Since its first outbreak in 1967 in Germany and Serbia, recurrent outbreaks have been reported in several African countries, with case fatality rates reaching up to 88% in some epidemics (2). Although MARV poses a serious public health risk and has epidemic potential, approved antiviral therapies specific to this virus are still lacking. This gap highlights the urgent need for effective and rational drug discovery (3).

The Marburg virus (MARV) genome encodes seven structural proteins. Among them, viral protein 35 (VP35), and the nucleoproteins (NP) play essential roles in viral replication, immune evasion, and virion assembly. VP35 (PDB ID: 4GH9, Crystal structure of Marburg virus VP35 RNA binding domain) functions as a polymerase cofactor and a potent interferon antagonist, thereby suppressing the host innate immune response and promoting viral replication. Whereas, NP (PDB ID: 4W2Q and 4W2O, Anti-Marburgvirus Nucleoprotein Single Domain Antibody C Complexed with Nucleoprotein C-terminal domain) encapsidates the viral RNA, which is critical for nucleocapsid formation and the regulation of viral transcription (4, 5). Owing to their essential roles in the viral life cycle, these proteins are considered promising molecular targets for antiviral intervention.

Structure-based drug discovery (SBDD) provides a computational framework for screening potential inhibitors against viral protein targets (6). Molecular docking predicts ligand-protein binding orientations and estimates binding affinities (7). whereas molecular dynamics (MD) simulations provide insights into conformational flexibility, interaction stability, and time-dependent structural changes under near-physiological conditions (8). In addition, binding free energy was estimated using the MM/GBSA approach to provide a more reliable assessment of interactions, incorporating solvation and entropic effects (9). Collectively, these computational strategies play an important role in antiviral drug discovery pipelines, especially for high-risk pathogens for which experimental investigations are restricted by biosafety requirements (10).

There have been a number of computational research pursuing potential inhibitors against the individual MARV proteins through docking-based virtual screenings. While promising molecules have been reported, a considerable number of studies have focused mainly on static docking results, lacking comprehensive dynamic validation and multi-target assessment. Considering the complex replication strategy of MARV and the coordinated roles of its structural proteins, a multi-target inhibition approach may offer greater therapeutic potential than single-target strategies (11). Incorporation of extended MD simulations and the free energy calculations can also contribute to increased robustness and predictive reliability of the computational analysis.

Natural bioactive compounds have been extensively studied as potential antiviral agents owing to their structural diversity, bioactivity, and traditional medicinal use (12). Withaferin A, a steroidal lactone derived from *Withania somnifera*, has demonstrated antiviral, anti-inflammatory, and immunomodulatory properties in various experimental contexts. Its pharmacological versatility and reported safety profile render it a suitable candidate for computational antiviral screening (13).

In this study, an integrated in silico study was conducted to examine the inhibitory potential of Withaferin A against three key proteins (VP35, two NP) of MARV. Their crystal structures were retrieved from the RCSB Protein Data Bank for subsequent computational analyses (14). Molecular docking was performed to predict binding affinities and interaction patterns, followed by 100 ns molecular dynamics simulations and MM/GBSA free energy analysis to evaluate dynamic stability and thermodynamically feasible binding. Additionally, pharmacoinformatics and toxicity profiling are also being performed in order to evaluate drug-likeness and safety parameters. Using a combination of multi-target docking, extended validation through MD simulations, and estimation of binding free energy in a unified computational approach, this work is aimed towards a comprehensive evaluation of Withaferin A as a potential lead compound against Marburg virus.

## 2. Materials and Methods

The stepwise methodology of the entire study is illustrated in **Figure 1**.

**Figure 1:**
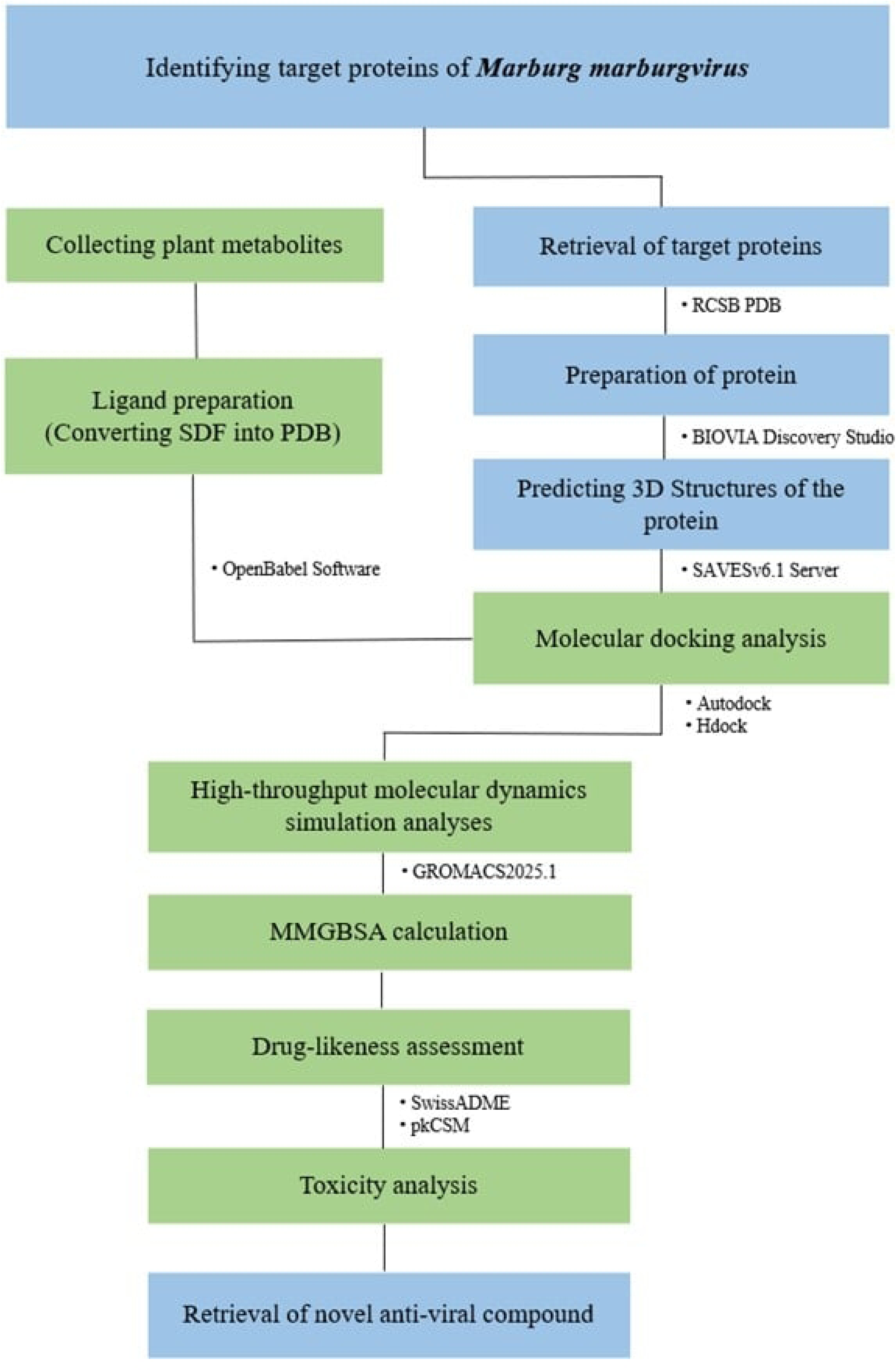
Schematic flowchart summarizing the methodological framework throughout the study.

### 2.1 Selection and Collection of Protein Sequences

The amino acid sequences of proteins VP35, and the nucleoproteins (NP) of Marburg virus were chosen. These proteins have been found to play vital roles in viral reproduction, immune response, and in forming a viral particle; therefore, they are a promising therapeutic target. Corresponding structural data were retrieved from the RCSB Protein Data Bank, ensuring the availability of experimentally resolved crystal structures appropriate for structure-based drug design (14).

### 2.2 Retrieval and Preparation of Target Proteins

The three-dimensional crystal structures of VP35 (PDB ID: 4GH9), and NP (PDB ID: 4W2Q and 4W2O) were downloaded from the RCSB Protein Data Bank (15). Protein preparation involved removal of crystallographic water molecules, co-crystallized ligands, and heteroatoms. Polar hydrogens were incorporated and Kollman charges were assigned using AutoDock tools. Energy minimization was performed to eliminate steric clashes for structural stability before docking.

### 2.3 Prediction of 3D Structures of the target Proteins

Since high-resolution crystal structures of the selected proteins were available in the Protein Data Bank, no additional homology modeling was required. Structural validation was performed using Ramachandran plot analysis to assess stereochemical quality and backbone dihedral angle distribution. Most residues have been found in favored and allowed regions, confirming the structural reliability for computational analysis.

### 2.4 Enlistment and Collection of Ligand Structure and Preparation

An initial library of 50 bioactive compounds was compiled based on reported antiviral and immunomodulatory properties. These compounds were subjected to preliminary virtual screening, from which Withaferin A (PubChem CID: 265237) was selected as the lead candidate based on its favorable binding performance and pharmacological relevance. The three-dimensional structure of Withaferin A was retrieved from the PubChem database (16). The downloaded SDF file was converted to PDB format using Open Babel (17). Energy minimization was conducted using the Universal Force Field (UFF) to obtain a stable, low-energy conformation suitable for subsequent docking analysis.

### 2.5 Molecular Docking

Molecular docking simulations were performed using AutoDock Vina (7). Grid boxes were configured to encompass the functional domains of VP35, and NP. The default exhaustiveness setting was used to ensure adequate conformational sampling. The resulting docked conformations were ranked based on predicted binding affinity values (kcal/mol). The lowest energy complexes were chosen for further analysis.

#### 2.5.1 Calculating the Binding Affinity of the Proteins and Ligands

AutoDock Vina’s scoring function was used for estimating the free energy of binding. Lower (more negative) binding energy values indicate more strongly predicted interactions between the ligand and protein. The most favorable docking conformation for each protein-ligand complex was identified based on its minimum binding energy and favorable interaction geometry.

#### 2.5.2 Study of the Preferred Binding Site of the Superior Metabolites

The preferred binding sites were analyzed by examining hydrogen bonding, hydrophobic interactions, electrostatic contacts, and van der Waals interactions using PyMOL and Discovery Studio Visualizer (18). Residues involved in ligand stabilization were identified, and their interactions within active or functionally relevant regions were further examined to clarify the possible inhibitory mechanism.

### 2.6 Molecular Dynamics Simulation Studies

The stability, flexibility, and dynamic properties of the protein and protein-ligand complexes were examined through molecular dynamics (MD) simulations. All simulations were conducted using GROMACS 2025.1 (19). The protein was parameterized with the CHARMM36 force field, while ligand topologies were generated using SwissParam (20). Prior to simulation, the systems were energy-minimized using a gradient-based algorithm for 2,500 steps to eliminate steric clashes and optimize structural geometry. The SPC water model was used to solvate each system in a cubic box with periodic boundaries. Na^+^ and Cl^-^ counter-ions were added to Gmx Genion to achieve overall charge neutrality.

After minimization, a two-stage equilibration was carried out. First, a 100 ps NVT equilibration was carried out using the v-rescale thermostat to stabilize the temperature at 310 K. This was followed by a 100 ps NPT equilibration using the Parrinello-Rahman barostat to maintain pressure at 1 bar and achieve density equilibration. Position restraints were applied to protein heavy atoms during equilibration. Bond constraints were imposed using the LINCS algorithm, and long-range electrostatic interactions were treated with the Particle Mesh Ewald (PME) method. After equilibration, three unrestrained 100 ns MD simulations were performed for all systems. Trajectories were saved at appropriate intervals for subsequent analyses.

#### 2.6.1 Root Mean Square Deviation (RMSD)

RMSD is a metric that quantifies the distance between frames. It is determined for every profile frame. The root mean square deviation of frame x is calculated.

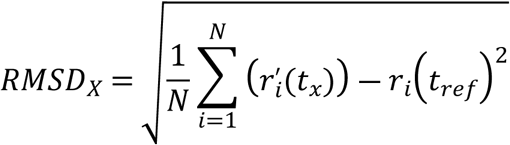

Where N is the number of selected atoms, r_i_ (t _ref_) represents the position of atom I in the reference structures denotes the position of the same atom in frame x after optimal structural superposition onto the reference. All RMSD calculations were performed using the gmx function in GROMACS2025.1, and the resulting trajectories were visualized using XmGrace.

#### 2.6.2 Root Mean Square Fluctuation (RMSF)

RMSF values can be used to determine the local flexibility based on residue displacements during the MD simulation. The Root Mean Square Fluctuation (RMSF) is useful for characterizing local changes along the protein chain. The RMSF for residue i is calculated using:

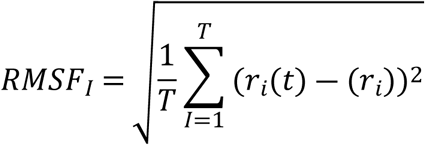

Where r_i_ (t) represents the position I at time t and r_i_ is its time averaged position. Unlike RMSD, which compares each frame to a reference structure, RMSF characterizes local fluctuations around the mean trajectory. Peaks in the RMSF profile correspond to flexible regions-typically loops and the N or C terminal segments, while lower values indicate more rigid, structurally stable regions of the protein.

#### 2.6.3 Radius of Gyration (Rg)

The radius of gyration (Rg) is the mass-weighted root mean square distance of a group of atoms around a common center of mass. Rg is widely used to evaluate the global compactness and folding stability of biomolecular systems during MD simulations. A stable Rg profile indicates maintenance of tertiary structure, whereas significant deviations imply expansion, unfolding, or structural reorganization. In this study, Rg was calculated for each trajectory frame using GROMACS analysis tools to monitor the compactness of the protein and protein-ligand complexes over the 100 ns simulation.

#### 2.6.4 Solvent Accessible Surface Area (SASA)

The Solvent Accessible Surface Area (SASA) was computed using the gmx sasa module in GROMACS 2025.1 along the 100 ns MD trajectory. SASA provides insights into the extent of solvent exposure and potential structural rearrangements at the protein surface. A default probe radius of 0.14 nm was applied, corresponding to the radius of a water molecule. SASA values were extracted at regular intervals for the entire protein-ligand complex to evaluate dynamic changes in surface exposure, folding stability, and solvent interactions throughout the simulation.

#### 2.6.5 Hydrogen Bond Analysis

The stability and formation of hydrogen bonds were measured on the 100 ns simulation in the gmx hbond module of GROMACS version 2025.1. Hydrogen bonds were identified using standard geometric criteria: a donor-acceptor distance ≤ 0.35 nm and a donor-hydrogen-acceptor angle ≥ 120°. The number of hydrogen bonds for each frame of the trajectory was also calculated to examine the intramolecular stability as well as to determine the effect of the ligand binding to the stability and distribution of the hydrogen bonds within the protein structure.

#### 2.6.6 MMGBSA Binding Free Energy

Molecular Mechanics/Generalized Born Surface Area (MM/GBSA) calculations were used to determine the free binding energy between the ligand and the protein and were done with gmx_MMPBSA tool (v1.6.3), which connects GROMACS trajectories to the AMBER free-energy calculations (21). This method breaks down the binding free energy (ΔG_bind) into molecular mechanical energy terms (van der Waals and electrostatic interactions) and solvation free-energy contributions (polar and nonpolar), according to the following equation:

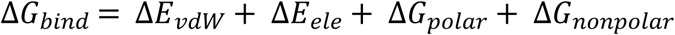

The Generalized Born (GB) implicit solvent model was used to compute the polar solvation energy, with the igb = 5 parameter corresponding to the GB-Neck2 model. The nonpolar solvation term (ΔG_nonpolar) was estimated from the solvent-accessible surface area using the LCPO method. Entropic contributions had been ignored, as is usual with comparative binding affinity studies, and has no effect on the relative ranking of ligands.

The 1001 snapshots were obtained in an equal manner in the 100 ns production trajectory (MD_center.xtc). MM/GBSA calculations were performed on the complex, the receptor (protein only) and the ligand, separately. The binding free energy was calculated using the standard thermodynamic cycle:

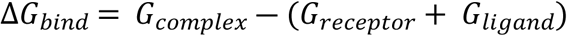

This protocol facilitated fine breakdown of contributions to binding including van der Waals, electrostatic, polar and nonpolar solvation contributions which offers a sound energetic assessment of the interactions between ligands and proteins.

### 2.7 Pharmacoinformatics Studies

SwissADME was used to predict the drug-likeness and pharmacokinetic properties of Withaferin A (22). Parameters analyzed included Lipinski’s Rule of Five compliance, molecular mass, hydrogen bond donors and acceptors, lipophilicity, and gastrointestinal absorption.

Additional pharmacokinetic predictions, including cytochrome P450 interactions and ADME parameters, were evaluated using pkCSM (23).

### 2.8 Toxicity Analysis

The pkCSM web server was used to determine the toxicity prediction of Withaferin A. Parameters evaluated included hepatotoxicity, AMES mutagenicity, cardiotoxicity (hERG inhibition), and acute toxicity prediction. These computational toxicity assessments provide preliminary safety insights before experimental validation.

## 3. Results

### 3.1 Retrieval of Targeted Proteins

The three-dimensional crystal structures of the Marburg virus proteins were successfully retrieved from the RCSB Protein Data Bank. The selected targets included viral protein 35 (VP35; PDB ID: 4GH9), and two nucleoproteins (NP; PDB ID: 4W2Q and 4W2O). All structures were downloaded in PDB format and prepared for subsequent computational analyses. Structural preprocessing involved the removal of heteroatoms and crystallographic water molecules, followed by the addition of hydrogen atoms and the assignment of appropriate charges. The prepared protein structures were then successfully utilized for downstream molecular docking and molecular dynamics simulations.

### 3.2 Molecular Modelling and Quality Assessment

The retrieved protein structures were evaluated for stereochemical quality using Ramachandran plot analysis. Then their 3D structures, Ramachandran plot and Errat quality values of the proteins are given in **Figure 2, Figure 3 and Figure 4**.

**Table 1.**
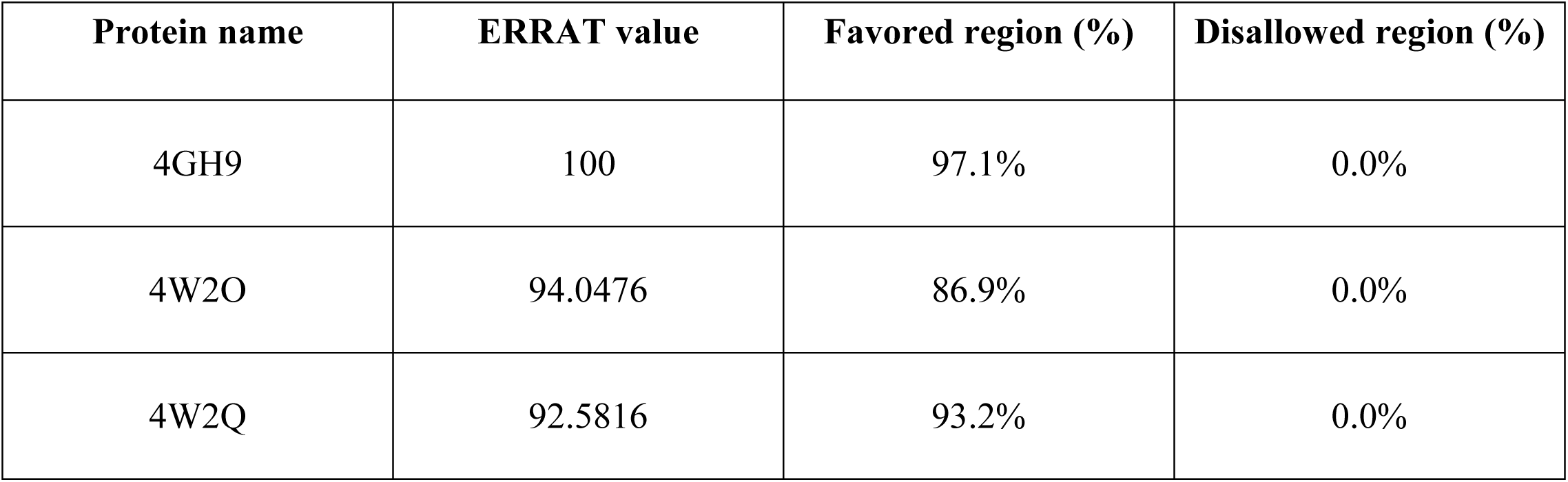
Refined protein model with their ERRAT values, Ramachandran plot results, and binding site residues.

| Protein name | ERRAT value | Favored region (%) | Disallowed region (%) |
| --- | --- | --- | --- |
| 4GH9 | 100 | 97.1% | 0.0% |
| 4W2O | 94.0476 | 86.9% | 0.0% |
| 4W2Q | 92.5816 | 93.2% | 0.0% |

**Figure 2:**
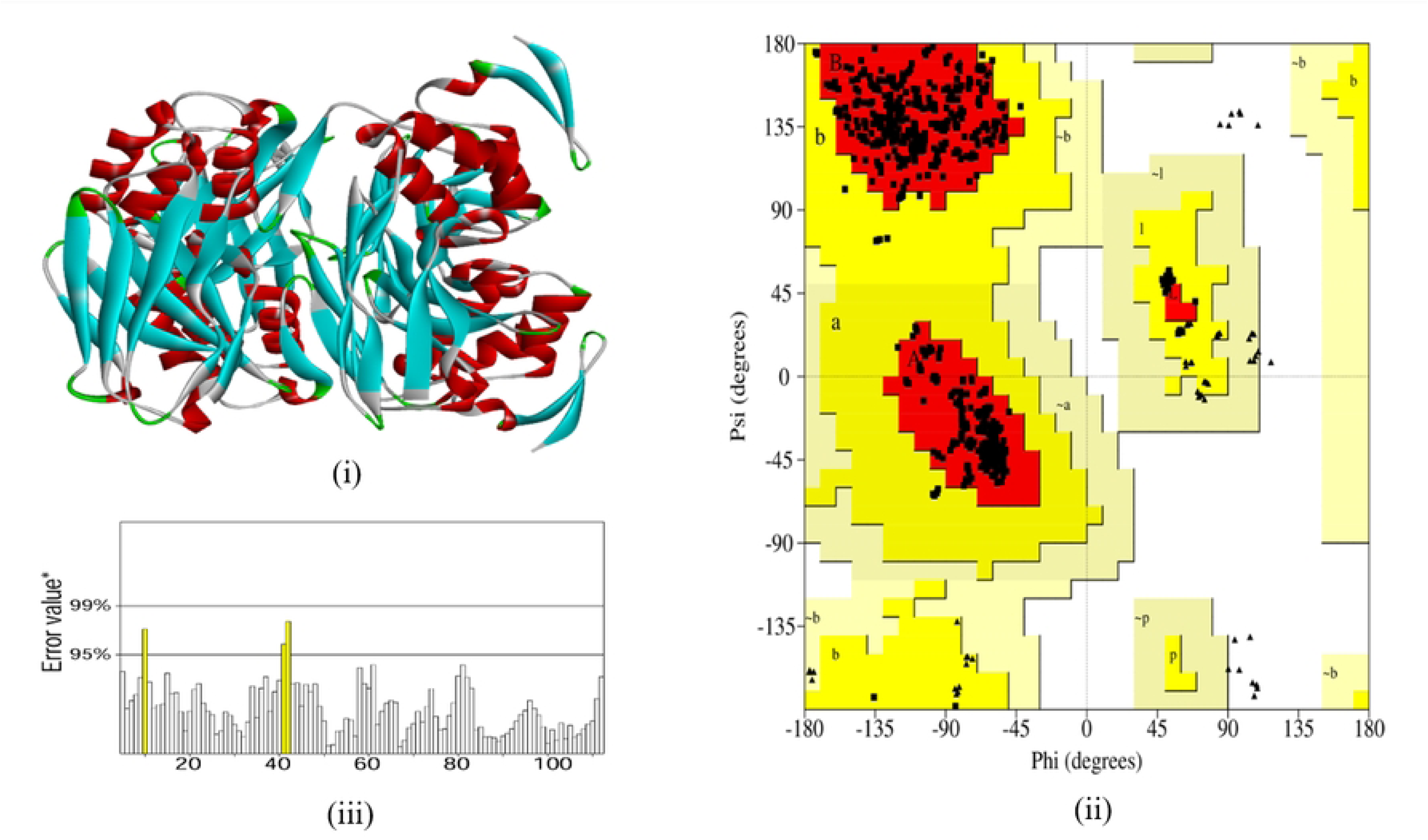
Structure prediction and validation of 4W2Q protein (i) 3D model, (ii) Ramachandran plot and (iii) Errat quality value.

**Figure 3:**
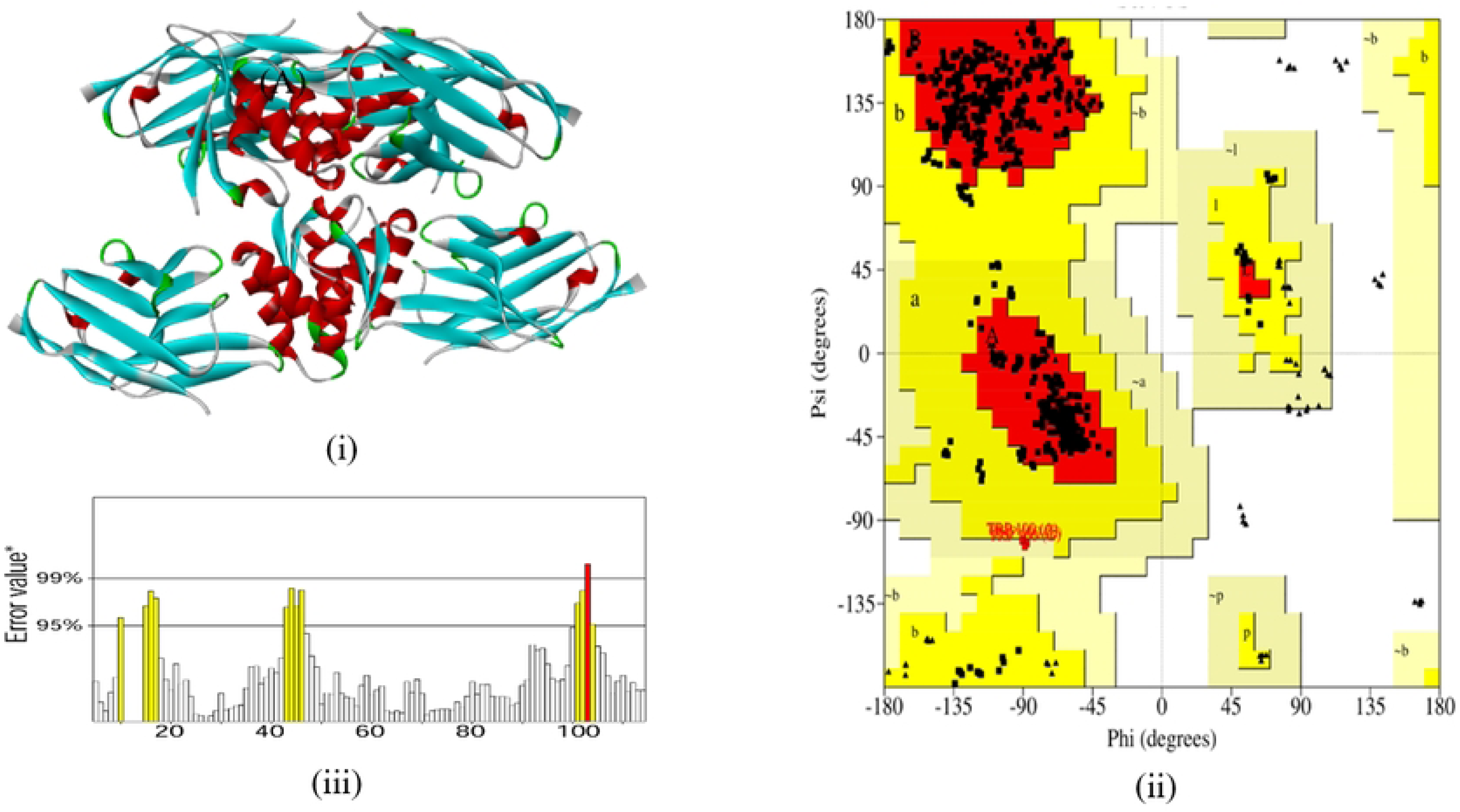
Structure prediction and validation of 4W2O protein (i) 3D model, (ii) Ramachandran plot and (iii) Errat quality value.

**Figure 4:**
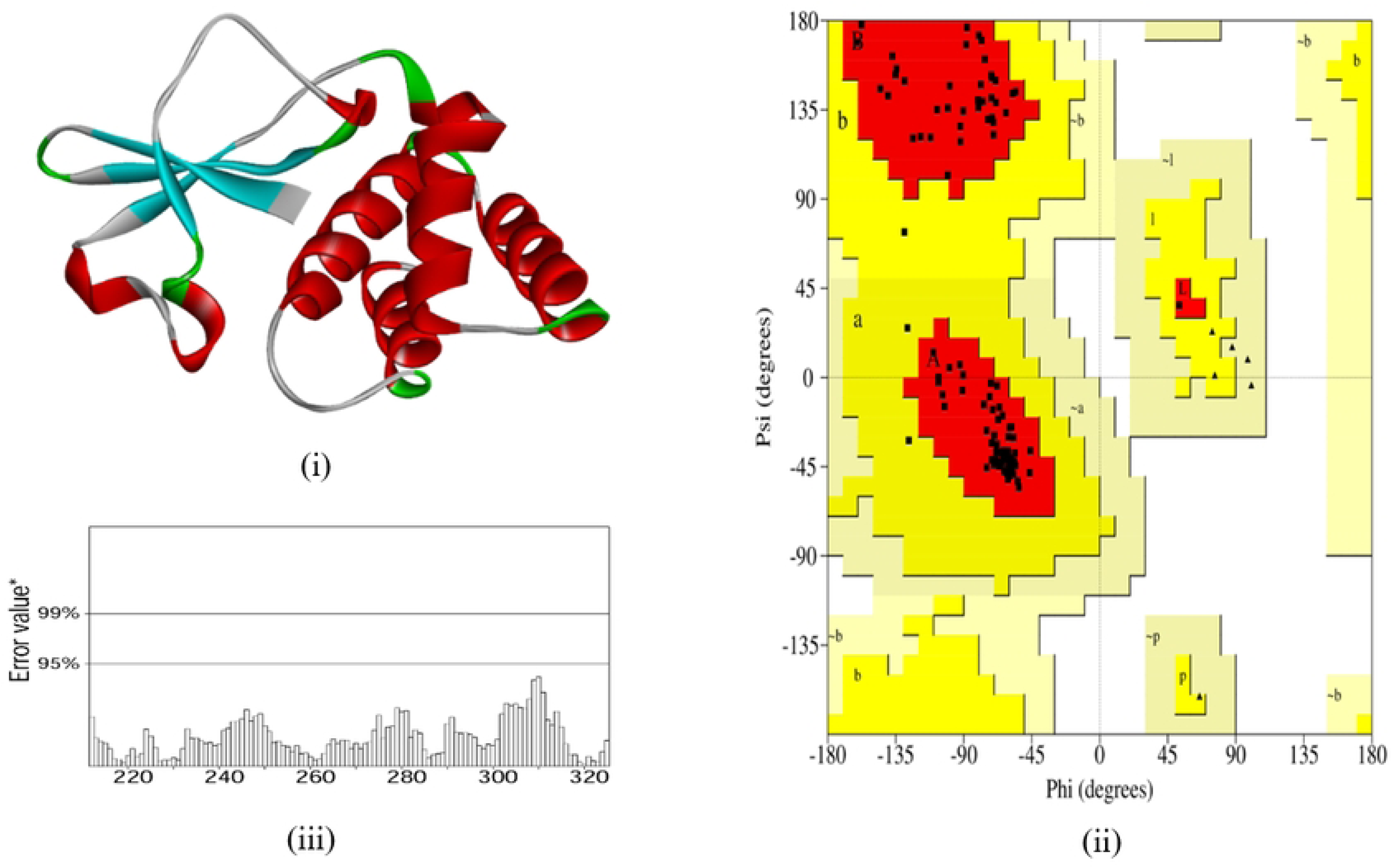
Structure prediction and validation of 4GH9 protein (i) 3D model, (ii) Ramachandran plot and (iii) Errat quality value.

The Ramachandran plots illustrated that most amino acid residues were found in favored and additionally allowed regions, with a minimal percentage of the residues found in disallowed regions. These findings verify the structural reliability and geometric stability of the chosen proteins for structure-based computational studies.

### 3.3 Molecular Docking and Binding Site Analysis

Molecular docking of Withaferin A (CID: 265237) with VP35 and NP of the Marburg virus, conducted using AutoDock Vina, is given in **Figure 5**. The predicted binding affinities demonstrated favorable interactions with all three targets. The docking scores were -8.2 kcal/mol for VP35 (PDB ID: 4GH9), -8.3 kcal/mol for NP (PDB ID: 4W2O), and -9.5 kcal/mol for NP (PDB ID: 4W2Q). Among the selected proteins, nucleoprotein (NP) exhibited the strongest binding affinity with Withaferin A, indicating a comparatively more stable ligand-protein interaction.

**Table 2.** Binding sites of the best metabolites with target protein.

| Protein | Ligand | HDock |  | CB-Dock | Hydrogen Bond Interactions | Hydrophobic Interactions (Alkyl / Pi-Alkyl) |
| --- | --- | --- | --- | --- | --- | --- |
|  |  | Docking Score | Confidence Score | Vina Score |  |  |
| 4W2O | Withaferin A | -201.33 | 0.7363 | -8.3 kcal/mol | LYS-145 (B), ASP-53 (A) | ALA-52 (A), LYS-475 (F) |
| 4GH9 | Withaferin A | -128.07 | 0.3921 | -8.2 kcal/mol | GLN-233, LYS-211 | LEU-215, PHE-218, ALA-214, LYS-237, ALA-210, TYR-240, LYS-241 |
| 4W2Q | Withaferin A | -184.40 | 0.6655 | -9.5 kcal/mol | GLU-649 (H), THR-551 (G) | LYS-314 (D), ALA-318 (D) |

**Figure 5:**
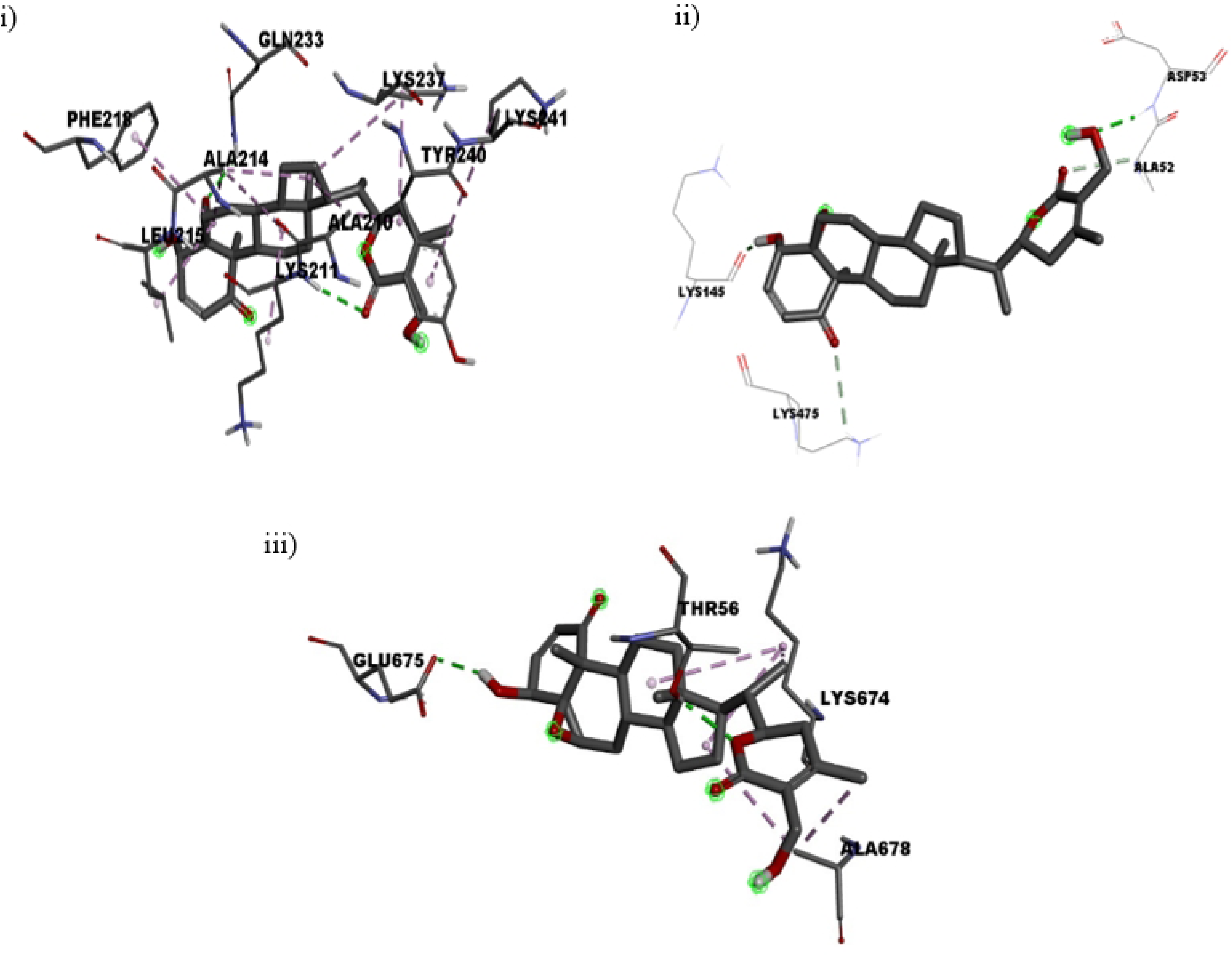
Docking conformation of Withaferin A with i. VP35 (4GH9), ii. NP (4W2O), iii. NP (4W2Q).

Among the three targets, the NP-Withaferin A complex exhibited the lowest binding energy.

Interaction analysis revealed that the formation of hydrogen bonds, hydrophobic interactions, and van der Waals contacts have been formed in the binding pockets of the proteins.

The 2D interaction diagrams of the proteins with Withaferin A are illustrated in **Figure 6**.

**Figure 6:**
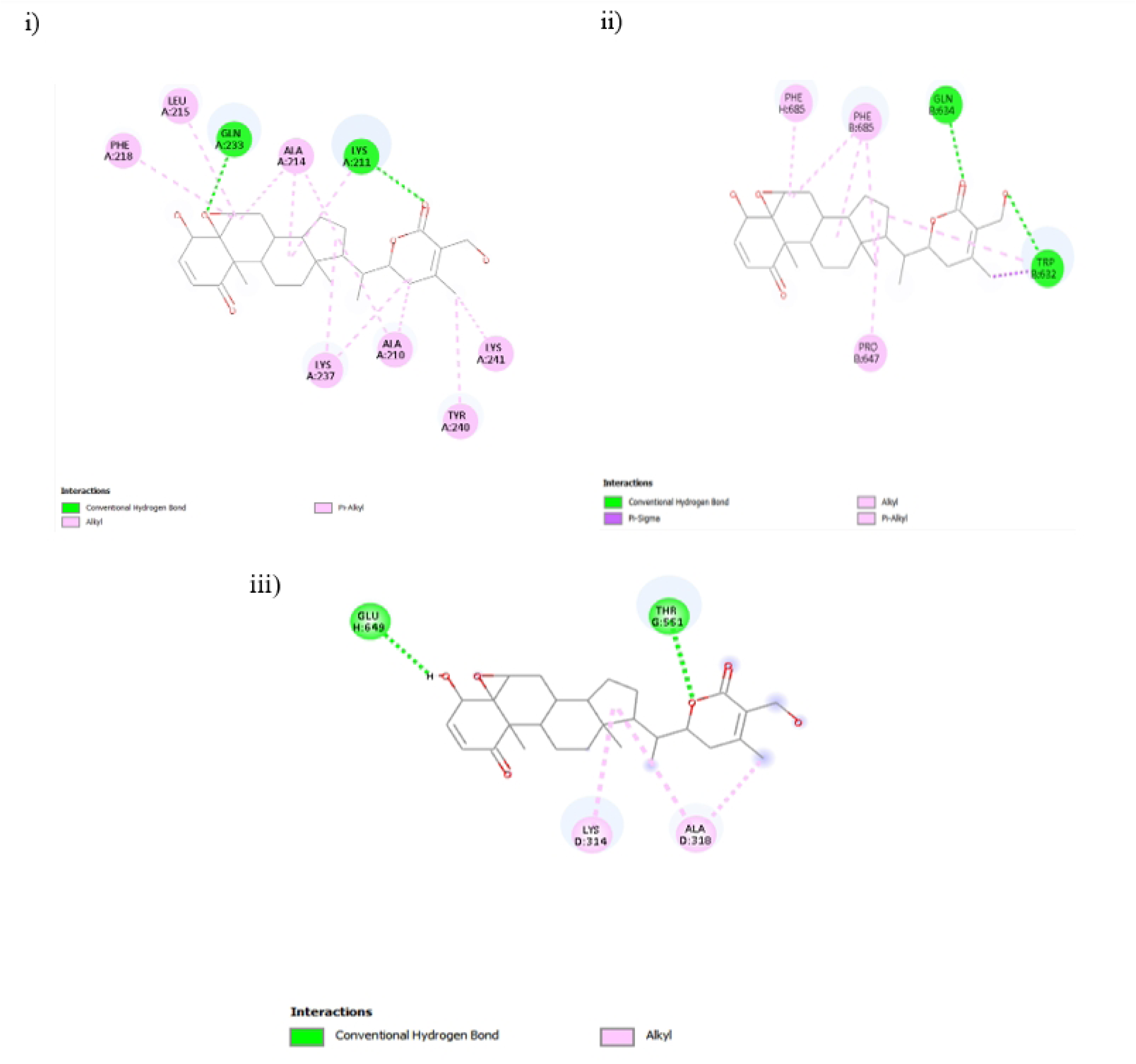
2D interaction diagram of i. VP35 (4GH9), ii. NP (4W2O), iii. NP (4W2Q) with Withaferin A complex.

The functional domains of each protein were found to have key interacting residues, which showed stable accommodation of their ligands in the predicted binding sites.

### 3.4 Analysis of Molecular Dynamics Simulation and MM/GBSA

#### 3.4.1 Root Mean Square Deviation (RMSD)

The RMSD profiles of the three complexes 4GH9-CID 265237 (WITHAFERIN A, black), 4W2Q-CID 265237 (WITHAFERIN A, red), and 4W2O-CID 265237 (WITHAFERIN A, green) demonstrated overall structural stability throughout the 100 ns simulation are given in **Figure 7**. The 4GH9 complex maintained RMSD values predominantly between 0.04-0.06 nm, with occasional spikes reaching approximately 0.18-0.19 nm around mid-simulation. The 4W2Q complex showed slightly lower deviation, fluctuating within 0.04-0.07 nm, with rare peaks near 0.16 nm. In contrast, 4W2O exhibited comparatively higher fluctuations, generally ranging between 0.06-0.10 nm, but without prolonged instability. The consistently low RMSD values (<0.2 nm) for all systems indicate stable protein-ligand conformations, with 4GH9 and 4W2Q showing comparatively better structural rigidity than 4W2O.

**Figure 7:**
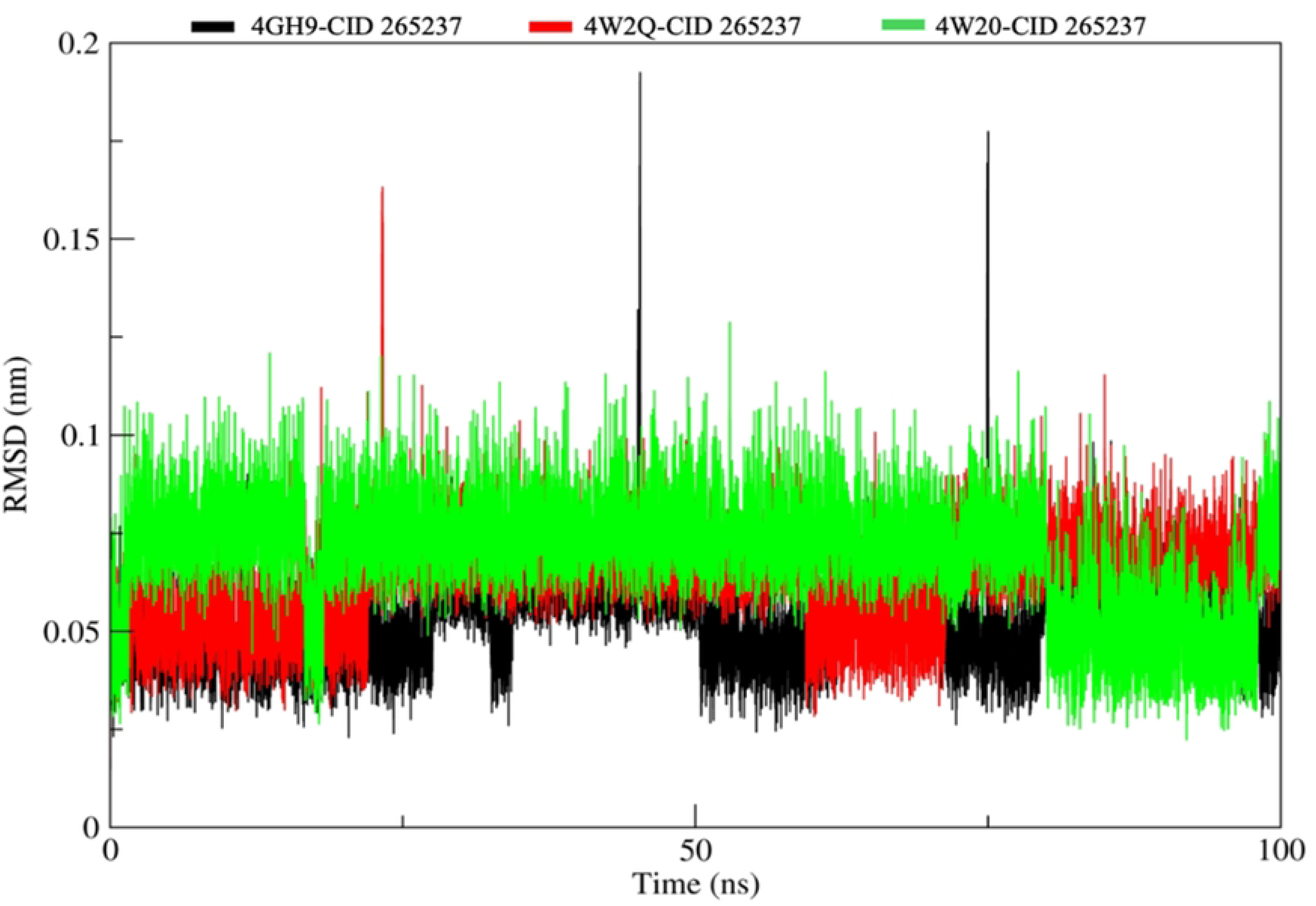
Time-dependent backbone root mean square deviation (RMSD) of the 4GH9-CID 265237, 4W2Q-CID 265237, and 4W2O-CID 265237 complexes during 100 ns of MD simulation.

#### 3.4.2 Root Mean Square Fluctuation (RMSF)

The RMSF analysis revealed residue-level flexibility differences among the complexes. The Residue-wise root means square fluctuation (RMSF) profiles of the three complexes during the molecular dynamic simulation are presented in **Figure 8**. For 4GH9, fluctuations were mostly confined to 0.05-0.10 nm, with a notable peak reaching approximately 0.20 nm near residue ∼310, indicating localized loop flexibility. The 4W2Q complex showed fluctuations primarily within 0.04-0.15 nm, with occasional peaks around 0.18-0.19 nm in the N-terminal region. Similarly, 4W2O exhibited RMSF values largely between 0.04-0.17 nm, with one prominent peak near 0.20 nm around residue ∼100. Overall, most residues in all systems remained below 0.10 nm, suggesting stable backbone dynamics, while higher peaks correspond to flexible loop regions rather than structural destabilization.

**Figure 8:**
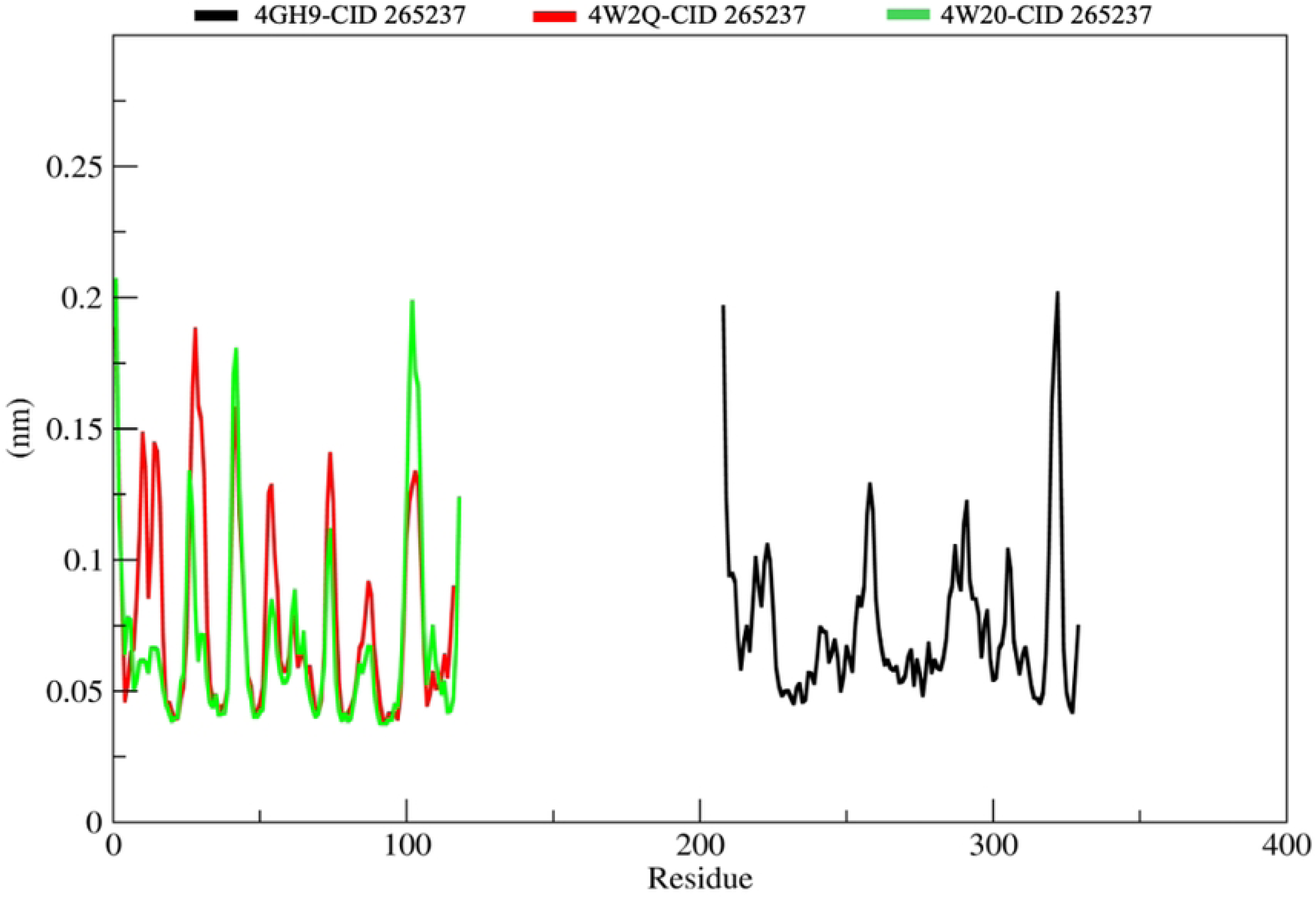
Residue-wise root means square fluctuation (RMSF) profiles of the three complexes during the molecular dynamic simulation.

#### 3.4.3 Radius of Gyration (Rg)

The radius of gyration analysis indicated compact structural behavior for all complexes across 100 ns (0-100,000 ps) is given in **Figure 9**. The 4GH9 complex maintained an average Rg of approximately 1.45-1.48 nm, showing minor fluctuations (∼±0.02 nm), suggesting a stable and compact structure. Both 4W2Q and 4W2O complexes exhibited slightly lower Rg values, averaging around 1.38-1.40 nm, with minimal deviation during the simulation. No significant expansion or unfolding events were observed in any system. The relatively constant Rg values confirm that ligand binding did not induce major conformational expansion, and the complexes remained structurally compact throughout the trajectory.

**Figure 9:**
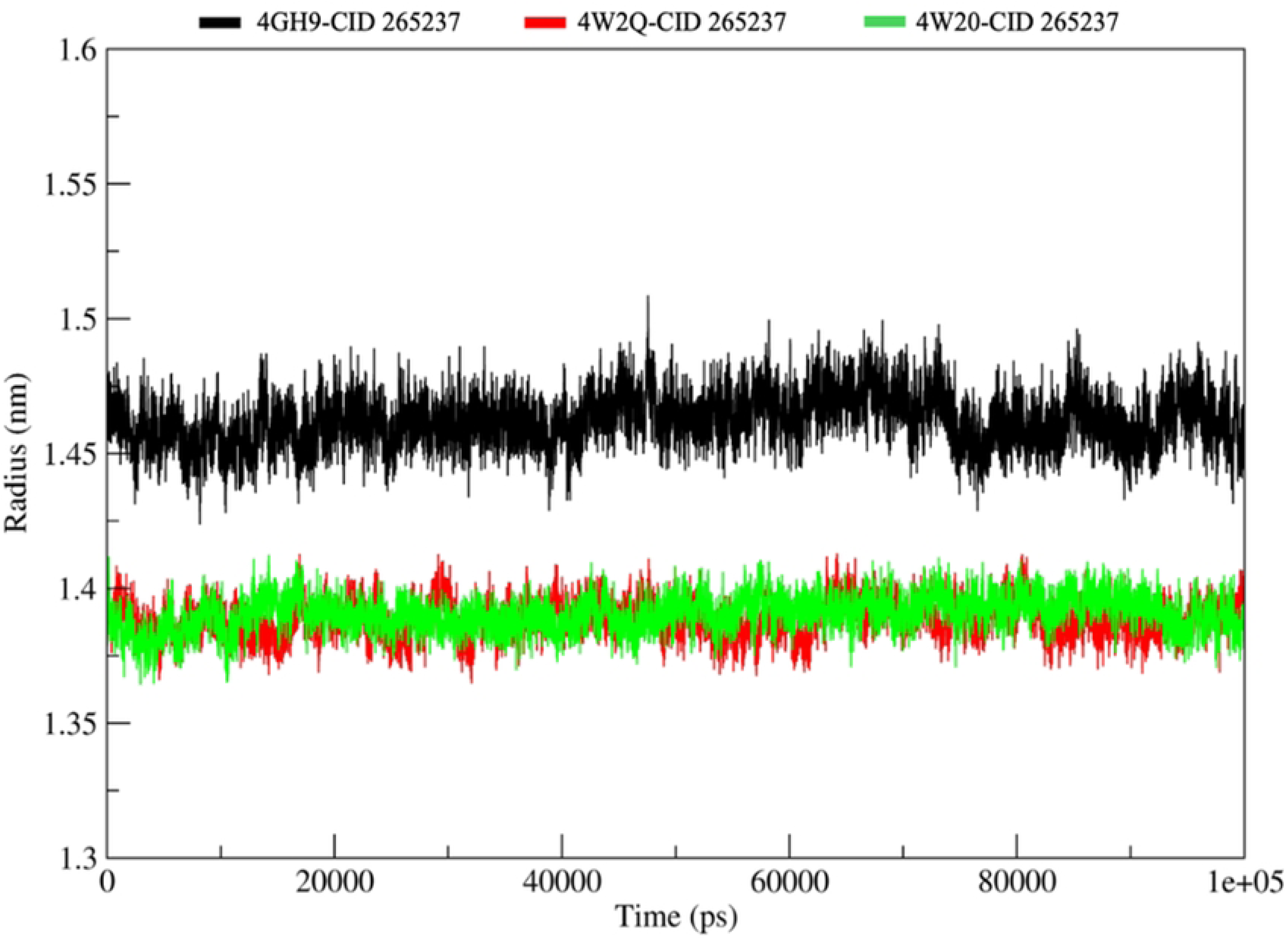
Radius of gyration (Rg) profiles of the three protein-ligand complexes over 100 ns of MD simulation.

#### 3.4.4 Solvent Accessible Surface Area (SASA)

The SASA profiles demonstrated moderate surface exposure variations among the complexes. Time-dependent solvent-accessible surface area (SASA) of the three protein-ligand complexes over the 100 ns molecular dynamics simulation, which are given in **Figure 10**. The 4GH9-CID 265237 (WITHAFERIN A) system showed SASA values fluctuating between 71-76 nm², indicating comparatively higher solvent exposure. The 4W2Q complex ranged between 68-74 nm², whereas 4W2O displayed slightly lower values, mostly between 64-70 nm². Although minor oscillations were observed throughout the simulation, no abrupt or sustained increases were detected, suggesting the absence of unfolding events. The relatively lower SASA in 4W2O corresponds with its slightly lower compactness variation observed in Rg analysis.

**Figure 10:**
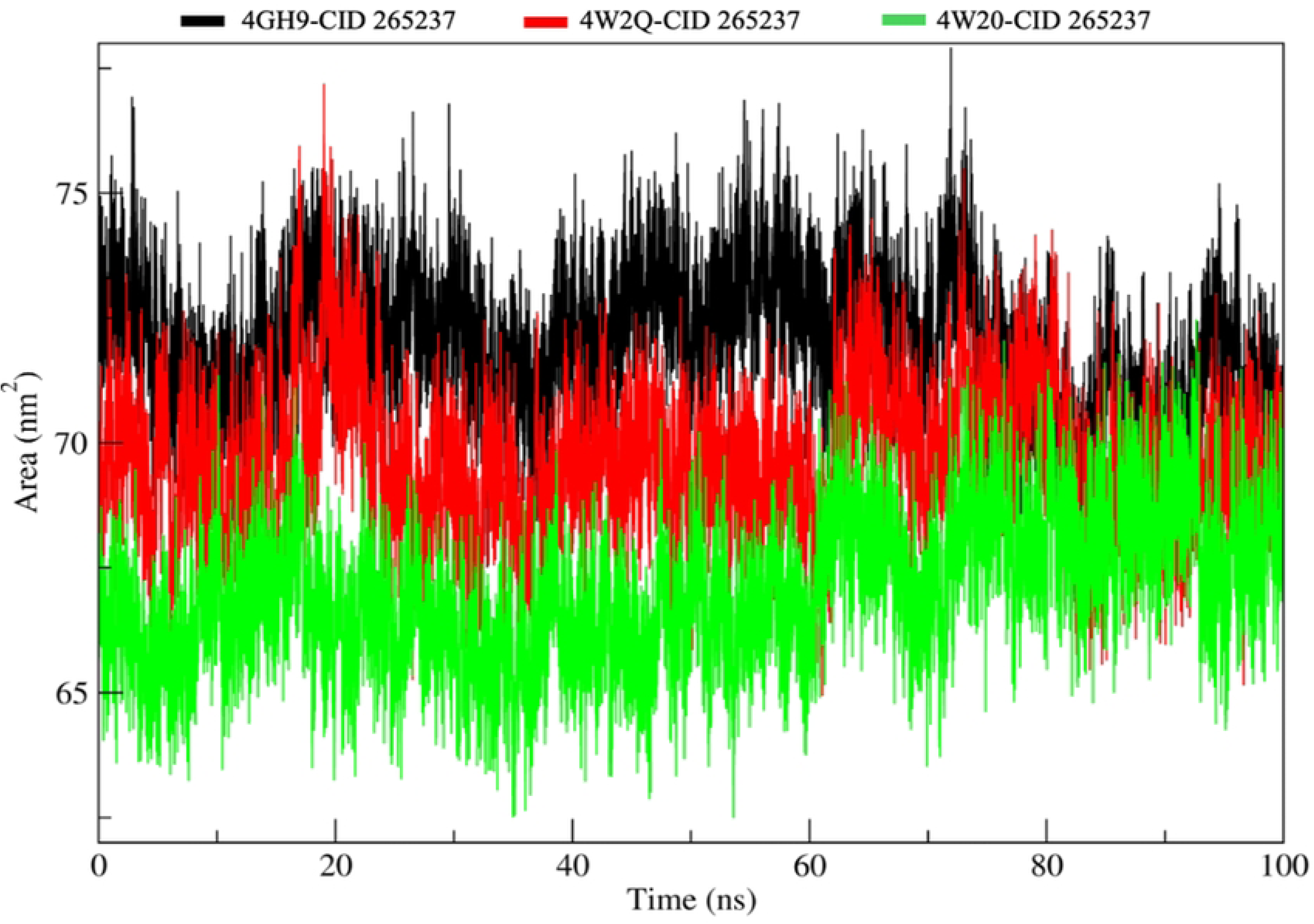
Time-dependent solvent-accessible surface area (SASA) of the three protein-ligand complexes over the 100 ns molecular dynamics simulation.

#### 3.4.5 Hydrogen Bonds

The hydrogen bond profiles for the three protein-ligand complexes during the 100 ns molecular dynamics simulation are shown in **Figure 11**. The 4GH9 complex formed between 0-3 hydrogen bonds, frequently maintaining 1-2 bonds during most of the trajectory. The 4W2Q complex also exhibited 0-3 hydrogen bonds, with sustained interactions particularly between 50-80 ns. Similarly, 4W2O maintained 0-3 hydrogen bonds, predominantly stabilizing at 1-2 bonds in the latter half of the simulation. Although transient bond breaking and reforming were observed, the consistent presence of hydrogen bonds across all systems supports stable ligand binding. Among them, 4W2Q showed comparatively more persistent hydrogen bonding during mid-to-late simulation phases.

**Figure 11:**
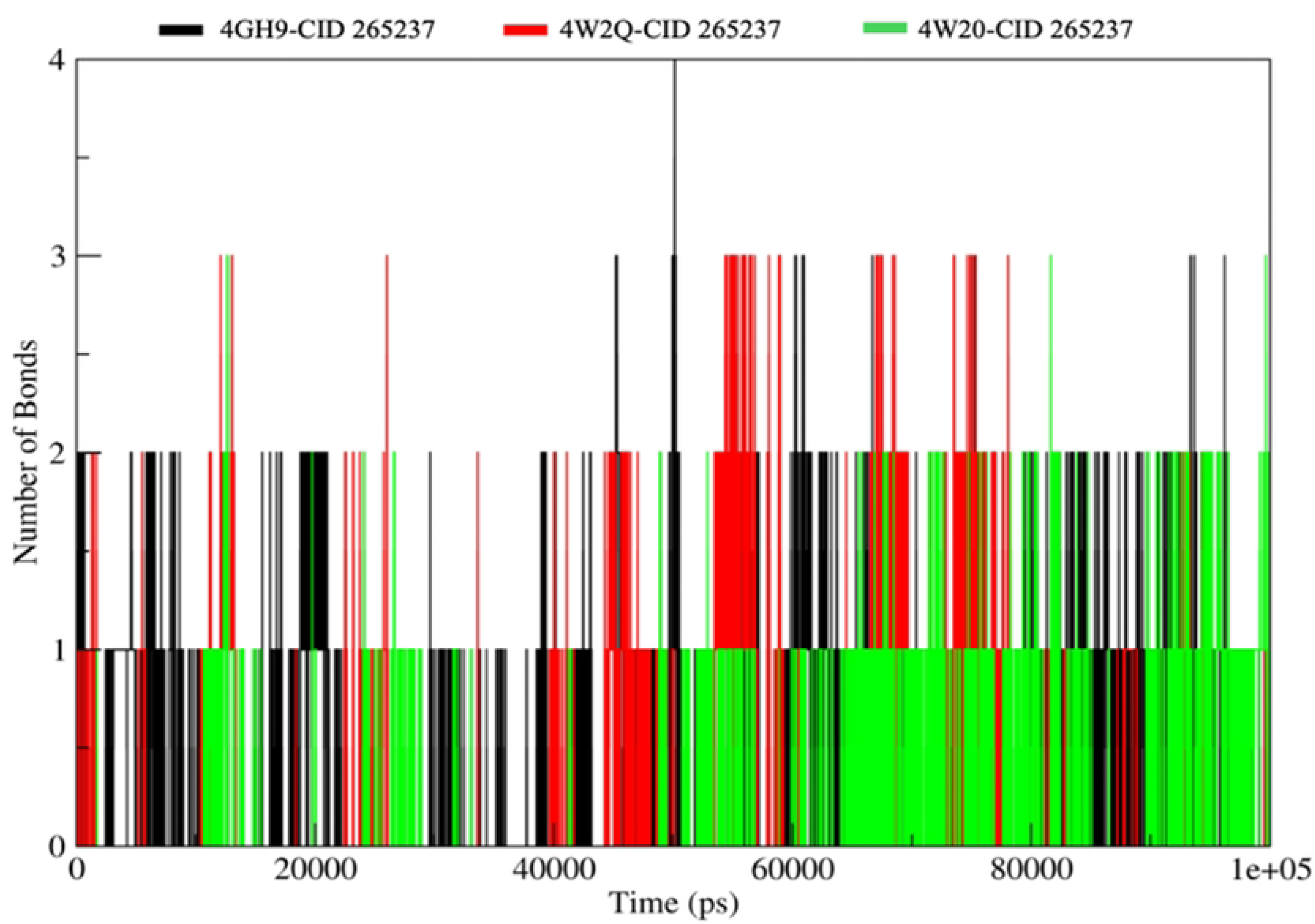
Hydrogen bond profiles for the three protein-ligand complexes during the 100 ns molecular dynamics simulation.

#### 3.4.6 MMGBSA

MM-GBSA binding free energy calculations for Withaferin A with VP35 (4GH9), NP (4W2O), and NP (4W2Q) of Marburg virus are given in **Figure 12**. Among the complexes, 4GH9-CID 265237 (WITHAFERIN A) exhibited the most favorable total binding free energy (ΔG = -6.99 kcal/mol), followed by 4W2O-CID 265237 (WITHAFERIN A) (ΔG = -5.68 kcal/mol), whereas 4W2Q-CID 265237 (WITHAFERIN A) showed the weakest binding affinity (ΔG = -1.16 kcal/mol). Negative ΔG values indicate favorable binding, with van der Waals interactions being the dominant contributor. The gas-phase interaction energy (GGAS) was strongly favorable for 4GH9 (-13.98 kcal/mol) and 4W2O (-11.95 kcal/mol), largely driven by significant van der Waals contributions (VDWAALS: -9.71 and -8.38 kcal/mol, respectively) and electrostatic interactions (EEL: -4.26 and -3.57 kcal/mol). In contrast, 4W2Q displayed substantially weaker van der Waals (-2.3 kcal/mol) and electrostatic (-1.03 kcal/mol) contributions, explaining its reduced binding affinity. Although the polar solvation energy (EGB) opposed binding in all systems (8.2, 7.35, and 2.47 kcal/mol for 4GH9, 4W2O, and 4W2Q, respectively), this unfavorable contribution was partially offset by favorable non-polar solvation energy (ESURF: -1.22, -1.09, and -0.3 kcal/mol). Overall, the stronger van der Waals and electrostatic interactions in 4GH9 and 4W2O compensated for solvation penalties, confirming more stable ligand binding compared to 4W2Q.

**Table 3:** MMGBSA Binding Energy Components.

| Energy Component | 4GH9-CID 265237<br>(WITHAFERIN A) | 4W2Q-CID 265237<br>(WITHAFERIN A) | 4W2O-CID 265237<br>(WITHAFERIN A) |
| --- | --- | --- | --- |
| VDWAALS | -9.71 | -2.3 | -8.38 |
| EEL | -4.26 | -1.03 | -3.57 |
| EGB | 8.2 | 2.47 | 7.35 |
| ESURF | -1.22 | -0.3 | -1.09 |
| GGAS | -13.98 | -3.33 | -11.95 |
| GSOLV | 6.98 | 2.17 | 6.27 |
| Total $\Delta G$ | -6.99 | -1.16 | -5.68 |

**Figure 12:**
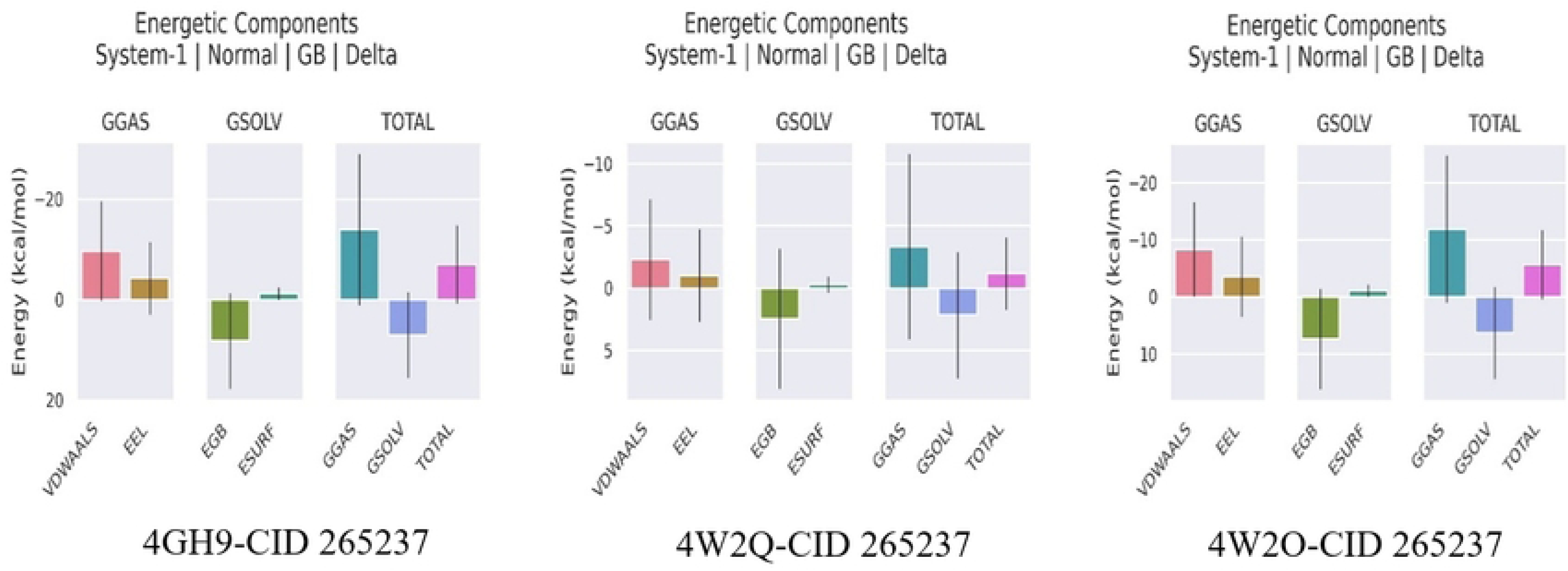
MM-GBSA binding free energy calculations for Withaferin A with VP35 (4GH9), NP (4W2O), and NP (4W2Q) of Marburg virus.

### 3.5 Pharmacoinformatics Studies

Drug-likeness and pharmacokinetic properties of Withaferin A were evaluated using SwissADME and pkCSM represented in **Figure 13**. The major drug-likeness parameters that Withaferin A met as expected were acceptable molecular weight, number of hydrogen bond donors and acceptors, and lipophilicity. Predicted gastrointestinal absorption was favorable, and cytochrome P450 interaction profiles were within acceptable ranges.

**Table 4.** ADME and drug-likeness properties of Withaferin A.

| Parameters |  | Withaferin A |
| --- | --- | --- |
| Physico-chemical parameters | Formula | C<sub>15</sub>H<sub>12</sub>O<sub>5</sub> |
|  | Molecular weight (g/mol) | 272.25 |
|  | Num. heavy atoms | 20 |
|  | Num. H-bond acceptors | 5 |
|  | Num. H-bond donors | 3 |
|  | Molar Refractivity | 71.57 |
|  | TPSA (Å<sup>2</sup>) | 86.99 |
| Lipophilicity | Log P (iLOGP) | 1.75 |
|  | Log P (XLOGP3) | 2.52 |
|  | Log P (WLOGP) | 2.19 |
|  | Log P (MLOGP) | 0.71 |
|  | Log P (SILICOS-IT) | 2.05 |
|  | Consensus Log P | 1.84 |
|  | Log P (iLOGP) | 1.75 |
| Pharmacokinetics | GI absorption | High |
|  | BBB permeant | No |
|  | P-gp substrate | Yes |
|  | CYP1A2 inhibitor | Yes |
|  | CYP2C19 inhibitor | No |
|  | CYP2C9 inhibitor | No |
|  | CYP2D6 inhibitor | No |
| **Water Solubility** | **Log S (ESOL)** | -3.49 |
| | Solubility (mg/ml; mol/l) | $8.74 \times 10^{-2}$ ; $3.21 \times 10^{-4}$ |
|  | Class | Soluble |
|  | **Log S (Ali)** | -3.99 |
| | Solubility (mg/ml; mol/l) | $2.77 \times 10^{-2}$ ; $1.02 \times 10^{-4}$ |
|  | Class | Soluble |
|  | **Log S (SILICOS-IT)** | -3.42 |
| | Solubility (mg/ml; mol/l) | $1.04 \times 10^{-1}$ ; $3.82 \times 10^{-4}$ |
|  | Class | Soluble |
| **Medicinal Chemistry** | PAINS | 0 |
|  | Brenk | 0 |
|  | Leadlikeness | Yes |
|  | Synthetic accessibility | 3.01 |

**Figure 13:**
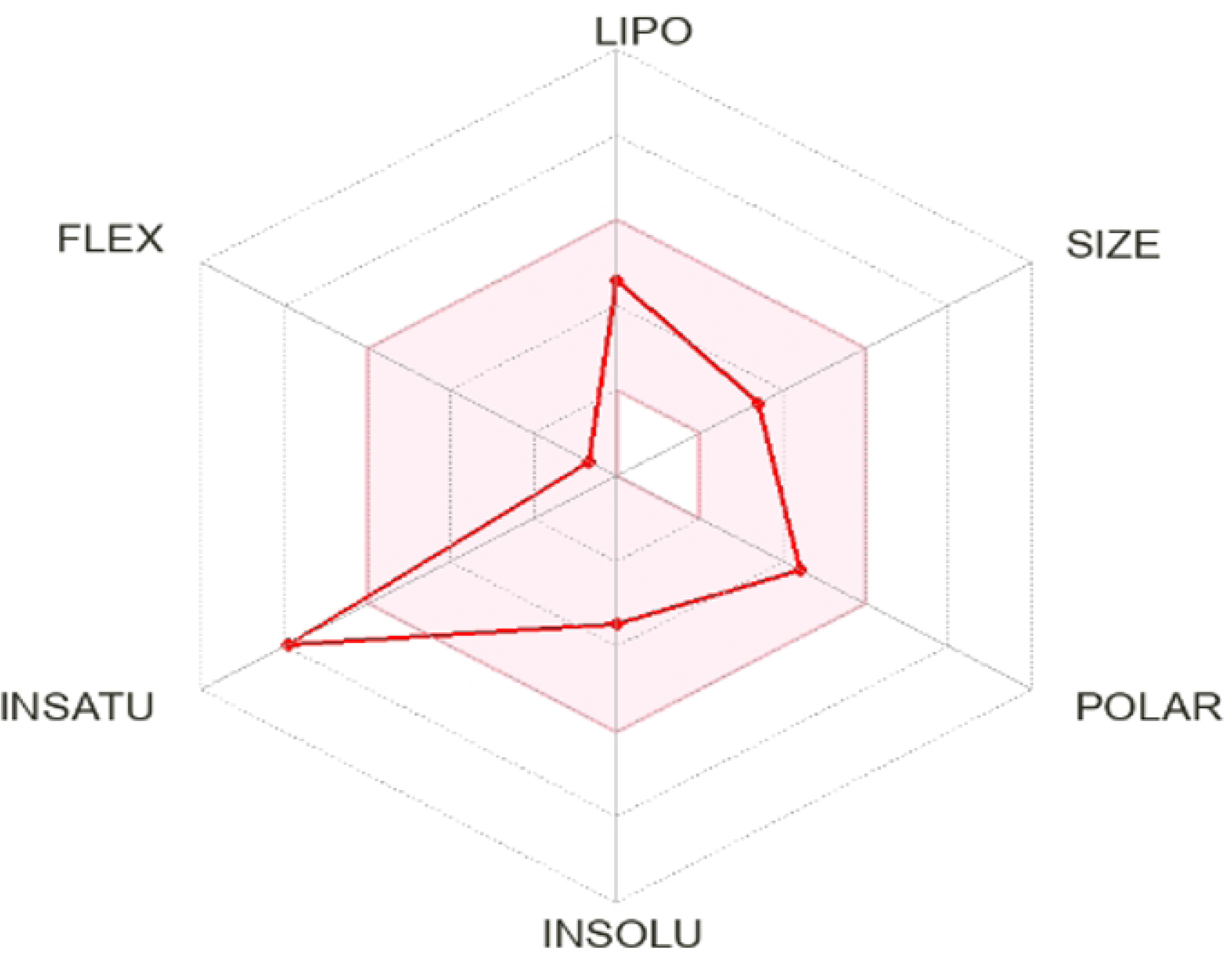
Pharmacokinetic properties of Withaferin A.

### 3.6 Toxicity Analysis of the Selected Metabolite

Toxicity prediction analysis was conducted using pkCSM. Withaferin A had been estimated to be non-mutagenic in AMES test and showed acceptable hepatotoxicity and cardiotoxicity profiles. Acute toxicity predictions were within permissible computational thresholds.

**Table 5.** Absorption, Distribution, Metabolism, Excretion and Toxicity parameter of selected metabolite.

| Category | Model Name | Predicted Value | Unit |
| --- | --- | --- | --- |
| Absorption | Water solubility | -3.224 | Numeric (log mol/L) |
| Absorption | Caco2 permeability | 1.029 | Numeric (log Papp in 10<sup>-6</sup> cm/s) |
| Absorption | Intestinal absorption (human) | 91.31 | Numeric (% Absorbed) |
| Absorption | Skin permeability | -2.742 | Numeric (log Kp) |
| Absorption | P-glycoprotein substrate | Yes | Categorical (Yes/No) |
| Absorption | P-glycoprotein I inhibitor | No | Categorical (Yes/No) |
| Absorption | P-glycoprotein II inhibitor | No | Categorical (Yes/No) |
| Distribution | VDss (human) | -0.015 | Numeric (log L/kg) |
| Distribution | Fraction unbound (human) | 0.064 | Numeric (Fu) |
| Distribution | BBB permeability | -0.578 | Numeric (log BB) |
| Distribution | CNS permeability | -2.215 | Numeric (log PS) |
| Metabolism | CYP2D6 substrate | No | Categorical (Yes/No) |
| Metabolism | CYP3A4 substrate | No | Categorical (Yes/No) |
| Metabolism | CYP1A2 inhibitor | Yes | Categorical (Yes/No) |
| Metabolism | CYP2C19 inhibitor | No | Categorical (Yes/No) |
| Metabolism | CYP2C9 inhibitor | No | Categorical (Yes/No) |
| Metabolism | CYP2D6 inhibitor | No | Categorical (Yes/No) |
| Metabolism | CYP3A4 inhibitor | No | Categorical (Yes/No) |
| Excretion | Total clearance | 0.06 | Numeric (log ml/min/kg) |
| Excretion | Renal OCT2 substrate | No | Categorical (Yes/No) |
| Toxicity | AMES toxicity | No | Categorical (Yes/No) |
| Toxicity | Max. tolerated dose (human) | -0.176 | Numeric (log mg/kg/day) |
| Toxicity | hERG I inhibitor | No | Categorical (Yes/No) |
| Toxicity | hERG II inhibitor | No | Categorical (Yes/No) |
| Toxicity | Oral rat acute toxicity (LD<sub>50</sub>) | 1.791 | Numeric (mol/kg) |
| Toxicity | Oral rat chronic toxicity<br>(LOAEL) | 1.944 | Numeric (log mg/kg_bw/day) |
| Toxicity | Hepatotoxicity | No | Categorical (Yes/No) |
| Toxicity | Skin sensitisation | No | Categorical (Yes/No) |
| Toxicity | *T. pyriformis* toxicity | 0.369 | Numeric (log µg/L) |
| Toxicity | Minnow toxicity | 2.136 | Numeric (log mM) |

## 4. Discussion

Previous computational studies that focused on Marburg virus (MARV) proteins have mostly used molecular docking approaches and virtual screening methods in order to discover potential inhibitory compounds. For example, several studies targeting VP35 inhibitors have been screened using natural product libraries, and reported acceptable docking scores for selected hits. However, subsequent analyses were often limited to interaction profiling and ADMET predictions, without performing extensive molecular dynamics validation (24). In a similar approach, repurposing studies on other MARV proteins have relied primarily on docking and pharmacokinetic evaluations to identify potential inhibitors. But, comprehensive long-timescale MD simulations were generally not employed to verify the stability of these interactions under dynamic conditions (25). These approaches are important for early-stage lead discovery. However, static docking snapshots overlook important factors such as protein flexibility, ligand-induced conformational changes, and solvent-mediated interactions, all of which are critical for molecular recognition and binding energetics (7).

A major shortcoming of many previous studies in which only docking scores were used for ranking candidate compounds. Docking-derived poses may represent energetically favorable conformations within a rigid environment. Yet, these conformations may prove unstable or transient when subjected to solvated, dynamic conditions that more accurately mimic the intracellular environment (26). In addition, several virtual screenings have focused on individual viral targets, which can limit the translational relevance of the identified compounds. Because the intricate and multi-protein nature of viral replication and immune evasion are complex (27). Although several amino acid-based interaction studies targeting MARV VP35 or the viral methyltransferase domain have reported promising binding affinities, these interactions were not consistently validated through extended MD simulations or time-resolved stability analyses (25).

In the current study, these shortcomings have been mitigated with a multi-target in silico evaluation of Withaferin A against three structurally and functionally diverse MARV proteins: the VP35, and nucleoproteins (NP). By integrating molecular docking with extended (100 ns) MD simulations and MM-GBSA binding free energy calculations, this current collaborative work offers a dynamic evaluation of the ligand-protein interactions beyond static docking predictions.

The MD analyses revealed stable backbone RMSD profiles, limited residue fluctuations in key binding regions, preserved structural compactness, and sustained intermolecular hydrogen bonding across the simulation trajectories. These results confirm that the docked complexes maintained dynamic stability under simulated physiological conditions. Together, these observations suggest that the ligand remained stably bound over time, rather than representing transient or non-physiological docking conformations.

The MM-GBSA binding free energy calculations offered a quantitative insight into the energetic contributions governing complex formation. Several complexes showed moderate binding free energies, largely driven by van der Waals and electrostatic interactions, whereas solvation penalties reduced the overall free energy gain. Notably, one protein-ligand complex exhibited substantially weaker binding energetics, indicating that Withaferin A does not interact uniformly across all MARV targets. This variability highlights the value of a multi-target evaluation and illustrates that structural stability observed in MD simulations does not always translate to strong binding affinity. Overall, these results suggest that Withaferin A exhibits measurable but moderate interaction strength, consistent with lead compounds rather than an inhibitor with relatively high affinity.

In addition to interaction analyses, many previous MARV-focused computational studies have lacked comprehensive pharmacoinformatics and toxicity assessments, which are crucial for prioritizing candidates in early-stage. To address this, the present study incorporated in silico ADME and toxicity evaluations using SwissADME and pkCSM to evaluate drug-likeness, pharmacokinetics, and safety parameters. The results indicated acceptable drug-like properties and a favorable predicted toxicity profile, supporting the suitability of Withaferin A as an early-stage antiviral lead from a pharmacological perspective. Nevertheless, these predictions are still by nature computational and must be experimentally confirmed. The results showed good drug-like characteristics and predicted toxicity profile, demonstrating the suitability of Withaferin A as an early-stage antiviral lead from a pharmacological point of view. Nevertheless, these predictions are still by nature computational and must be experimentally confirmed.

To the best of our knowledge, no previous study has comprehensively evaluated the inhibitory potential of Withaferin A against multiple Marburg virus protein targets using an integrated docking, molecular dynamics simulation, and MM-GBSA approach. By overcoming key limitations of the previous studies, i.e., lack of dynamic validation, single-target focus, and incomplete pharmacoinformatics assessment, this study provides a more thorough and realistic evaluation of ligand-protein interactions. Although, the binding energies observed were moderate and suggest that Withaferin A may not act as a potent standalone inhibitor, it represents a promising scaffold for further structural optimization and experimental validation in the context of multi-target antiviral drug discovery against Marburg virus.

## 5. Conclusion

The present study used an integrated computational approach to assess the antiviral potential of Withaferin A (CID 265237) against three vital proteins of Marburg virus, including VP35 (4GH9), NP (4W2Q and 4W2O). Molecular docking studies demonstrated favorable binding affinities of Withaferin A toward all target proteins. These interactions illustrate its potential to engage functionally relevant domains associated with viral replication, immune evasion, and virion assembly.

To validate the docking predictions, 100 ns molecular dynamics simulations were conducted using GROMACS. The simulations exhibited stable RMSD convergence, controlled residue flexibility, sustained structural compactness, and persistent hydrogen bonding over the simulation timeframe. MM-GBSA binding free energy analysis further supported thermodynamically favorable interactions. The binding was predominantly driven by van der Waals forces, with electrostatic interactions contributing to overall complex stability. Such dynamic analyses offer greater assurance regarding the stability and persistence of the protein-ligand complexes beyond static docking predictions.

Importantly, the multi-target evaluation approach used in this study distinguishes it from previous single-target investigations. By exhibiting stable interactions with VP35, and NP; Withaferin A might have the potential to interfere with multiple stages of the viral life cycle, which could minimize the chances of developing resistance.

Although the results are based on computational predictions and require experimental validation. Nevertheless, the findings collectively highlight Withaferin A as a promising multi-target novel compound against Marburg virus. This study represents mechanistic insights and establishes a foundation for future research conducted both in vitro and in vivo, to develop effective antiviral therapeutics against MARV.

## 7. Acknowledgements

The authors sincerely acknowledge the contributions of all co-authors for their involvement in the conceptualization, design, analysis, and preparation of this manuscript. We also thank the Department of Animal and Fish Biotechnology, Faculty of Biotechnology and Genetic Engineering, Sylhet Agricultural University, for providing the necessary facilities and support to conduct this study.

